# Effect of intensity of agronomic practices on the yield of two breeding types of winter oilseed rape cultivars

**DOI:** 10.1101/2019.12.20.884304

**Authors:** Marek Wójtowicz, Andrzej Wójtowicz, Ewa Jajor, Marek Korbas, Franciszek Wielebski

**Affiliations:** Plant Breeding and Acclimatization Institute—National Research Institute, Independent Laboratory of Oilseed Crops Environmental Stress, Poznań, Poland; Plant Protection Institute—National Research Institute, Department of Mycology, Poznań, Poland

## Abstract

The effect of three fungicide treatment programmes and the level of spring nitrogen fertilisation on the seed yield of two types of cultivars of *Brassica napus* L. sown at two different seeding rates was studied in a field experiment carried out in a split-split-plot design. The subject of the study was an open-pollinated cultivar (Casoar) and a restored hybrid cultivar (Visby). Three plant protection programmes, two levels of spring nitrogen fertilisation (160 and 220 kg N·ha^-1^), and two different seeding rates for each cultivar (Visby—50 and 70 seeds·m^-2^; Casoar—60 and 80 seeds·m^-2^) were included. The most intensive protection programme comprised three fungicide treatments: first in autumn at the six-leaves-unfolded stage—BBCH 16, second in spring at the stem elongation stage—BBCH 33, and third at the full flowering stage—BBCH 65. One of two less intensive programmes of plant protection included fungicide application in autumn at the six-leaves-unfolded stage—BBCH 16 and at the full flowering stage—BBCH 65, while the second included fungicide application in spring at the stem elongation stage—BBCH 33 and at the full flowering stage—BBCH 65.

The effectiveness of the protection programmes and nitrogen fertilisation was influenced by the intensity of abiotic stress factors. However, the average yield from the plots protected against pathogens was significantly higher than that from the untreated plots. The increase of nitrogen fertilisation from 160 to 220 kg·ha^-1^ also caused significant increase of average seed yield. The yield of cultivar Visby was higher and less dependent on the seeding rate compared to cultivar Casoar. Cultivars responded similarly to plant protection programmes and the rate of nitrogen fertilisation in spring. Higher yields of Visby cultivar can be attributed to the higher number of seeds per silique and the higher number of siliques per m^2^.

## Introduction

Crop yields are shaped by advances in breeding progress, which result in varieties with favourable properties, and intensification of cultivation treatments. In the northern hemisphere characterised by a cold climate, the only commonly cultivated oilseed crop is oilseed rape (*Brassica napus* L.). The possibility of using oilseed rape in the food [1] and feed [25] industry, as well as in the generation of energy [26], especially biofuels, has significantly contributed to the increase in the cultivation area of this oilseed crop (0.6 million ha·year^-1^ between 1997 and 2017) [7]. Increase of the cultivation area as well as the level of yield of oilseed rape recorded since the early 1990s [9, 45] reflects the interest in rapeseed of the food and feed industry, which was gained after the introduction of the double-improved variety with no erucic acid and reduced glucosinolate, and the energy industry using rapeseed oil in biofuel production. The possibility of growing rapeseed for energy purposes has contributed to the increase in the cultivation area of this oilseed plant, and also to its higher yields obtained in Poland.

The sown area and yield of oilseed rape increased, respectively, from 0.4 million ha and 2.1 Mg·ha^-1^ on average during 2001–2003 before Poland’s accession to the European Union to 0.9 million ha and 3.0 Mg·ha^-1^ in the years 2014–2016 [32]. Such a significant increase in the area stimulates the demand for high- and stable-yielding cultivars that react efficiently to cultivation treatments and justifies the intensification of production strongly conditioned by economic factors. The yields of oilseed rape are shaped on the one hand by the yield potential of the variety [24], its tolerance/resistance to biotic and abiotic stress [41], and on the other hand by cultivation technology, the aim of which is to provide the most favourable conditions for the development of crops and to limit crop losses. The resistance of cultivars to biotic and abiotic stresses is particularly important. Cultivars that exhibit good tolerance to stressful conditions have a better chance of showing good yield potential because the seasons characterised by unfavourable weather conditions for the development of oilseed rape are not uncommon in Poland. For this reason, one of the objectives of the present research is to understand the adaptive abilities of the types of rapeseed cultivars studied to changing environmental conditions. The working hypothesis assumes the overriding role of cultivars’ resistance to abiotic stresses in shaping their yield level. The shortage of precipitation during the stages characterised by the greatest demand for water and low temperatures during winter dormant period, which can be seen frequently in Poland, affecting the development of plants may contribute to a significant decrease in the yield level of rape cultivars. In addition, excess of rainfall can limit the yields, especially in the case of cultivars susceptible to infection, because it induces infections and the spreading of pathogens. An effective way to protect plants from pathogens is fungicide treatment [12, 29, 47]. The dependence of humidity conditions of the oilseed rape canopy on the plant density derived from the amount of seeds sown justifies research into the effectiveness of protection against pathogens at different sowing rates. The constant increase of importance of chemical protection against pathogens in the technology of oilseed rape [42] and the decisive significance of the amount of seeds sown for the yield level confirm the accuracy of the selection of these experimental factors. Moreover, the variability of cultivars’ resistance to pathogens [14-15] justifies undertaking research aimed at determining their response to the intensification of this factor of production, although it is rare to find results presenting variation in the yields of cultivars resulting from the intensification of protection [10]. Another factor included in this research mainly due to its stimulating impact on the development and yield of rapeseed [2, 6, 30-31], as well as on infection by pathogens [37], is spring nitrogen fertilisation. Understanding the impact of the intensification of nitrogen fertilisation on the development and yielding of the types of rapeseed cultivars compared here is another goal of the present work. The choice of experimental factors studied in this work was also dictated by their significant role in determining cultivation costs. According to Budzyński and Ojczyk [5], fertilisation and protection account for almost 80% of the cultivation costs of oilseed rape.

## Materials and methods

The experiment was conducted in 2012–2014 at the Experimental Station of Plant Breeding Smolice Ltd, Co. in Łagiewniki (N 51° 46′, E 17° 14′). It was a three-factor type and was laid out in a split-split-plot design with four replications. The main plot factor was the programme of protection against pathogens (Table 1). Three programmes of fungicide application and a control treatment were included in the experiment. The most intensive protection programme comprised three fungicide applications: first in autumn at the six-leaves-unfolded stage—BBCH 16, second in spring at the stem elongation stage—BBCH 33, and third at the full flowering stage—BBCH 65. In addition, two less intensive programmes of plant protection were included: the first one involved fungicide application in autumn at the six-leaves-unfolded stage—BBCH 16 and at the full flowering stage—BBCH 65, while the second involved fungicide application in spring at the stem elongation stage—BBCH 33 and at full flowering stage—BBCH 65. In autumn, metconazole (5-[(4-chlorophenyl)methyl]-2,2-dimethyl-1-(1H-1,2,4-triazol-1-ylmethyl) cyclopentanol) was applied at 60 g·ha^-1^. At the stem elongation stage, prothioconazole (2-[2-(1-chlorocyclopropyl)-3-(2-chlorophenyl)-2-hydroxypropyl]-1,2-dihydro-3H-1,2,4-triazole-3-thione) was applied at 80 g·ha^-1^ and tebuconazole (α-(2-(4-chlorophenyl)ethyl)-α-(1,1-dimethylethyl)-1H-1,2,4-triazole-1-ethanol) was applied at 160 g·ha^-1^. At the flowering stage, dimoxystrobin ((αE)-2-[(2,5-dimethylphenoxy)methyl]-α- (methoxyimino)-N-methylbenzeneacetamide) at 100 g·ha^-1^ and boscalid (2-chloro-N-(4′-chloro[1,1′-biphenyl]-2-yl)-3-pyridinecarboxamide) at 100 g·ha^-1^ were applied. The subplot factor was the rate of nitrogen fertilisation. Nitrogen fertiliser was applied at two levels: 160 and 220 kg N·ha^-1^. The sub-subplot factor was represented by cultivars sown at different seeding rates; open-pollinated cultivar Casoar was sown at a seeding rate of 60 and 80 seeds·m^-2^, while the fertility-restored hybrid cultivar Visby was sown at 50 and 70 seeds·m^-2^. The experiment was carried out on proper brown soil formed from heavy clay sand, on light or middle clay, of quality class IIIa and good wheat complex. The winter oilseed rape crops previously cultivated on the soil were rye, lucerne, and spring wheat. The chemical constituents of the soil were as follows: P_2_O_5_, 221–276 mg·kg^-1^; K_2_O, 135–191 mg·kg^-1^; Mg, 31–73 mg·kg^-1^; and N, min 6.6–9.4 mg·kg^-1^. The pH of the soil ranged from 6.3 to 7.2 when measured using 1 M KCl. Before sowing, the field received 20–25 kg N·ha^-1^ (ammonium nitrate), 51–80 kg P_2_O_5_·ha^-1^ (triple superphosphate), and 105–112 kg K_2_O·ha^-1^ (60% potash salt). Winter oilseed rape was sown with 30-cm row spacing on 26–29 August. Plants on all investigated plots were protected using herbicides and insecticides. The dicotyledonous weeds were controlled with metazachlor (2-chloro-N-(2,6-dimethylphenyl)-N-(1H-pyrazol-1-ylmethyl)acetamide) at a dose of 999 g·ha^-1^ and quinmerac (7-chloro-3-methyl-8-quinolinecarboxylic acid) at a dose of 249 g·ha^-1^, and volunteer cereals with herbicide cycloxydim (2-[1-(ethoxyimino)butyl]-3-hydroxy-5- (tetrahydro-2H-thiopyran-3-yl)-2-cyclohexen-1-one) at a dose of 150 g·ha^-1^. Insects were controlled using lambda-cyhalothrin ((R)-cyano(3-phenoxyphenyl)methyl (1S,3S)-rel-3-((1Z)-2-chloro-3,3,3-trifluoro-1-propenyl)-2,2-dimethylcyclopropanecarboxylate, 6.25 g·ha^-1^), tiachloprid ((Z)-(3-((6-chloro-3-pyridinyl)methyl)-2-thiazolidinylidene)cyanamide, 6.0 g·ha^-1^), deltamethrin ((S)-cyano(3-phenoxyphenyl)methyl (1R,3R)-3-(2,2-dibromoethenyl)-2,2-dimethylcyclopropanecarboxylate, 60 g·ha^-1^), and acetamiprid ((1E)-N-[(6-chloro-3-pyridinyl)methyl]-N′-cyano-N-methylethanimidamide, 24 g·ha^-1^). Maturated plants were harvested without swathing using a small-plot combine harvester on 13, 23, and 15 July in 2012, 2013, and 2014, respectively. The plot area to be harvested was 9.6 m^2^.

**Table 1.**
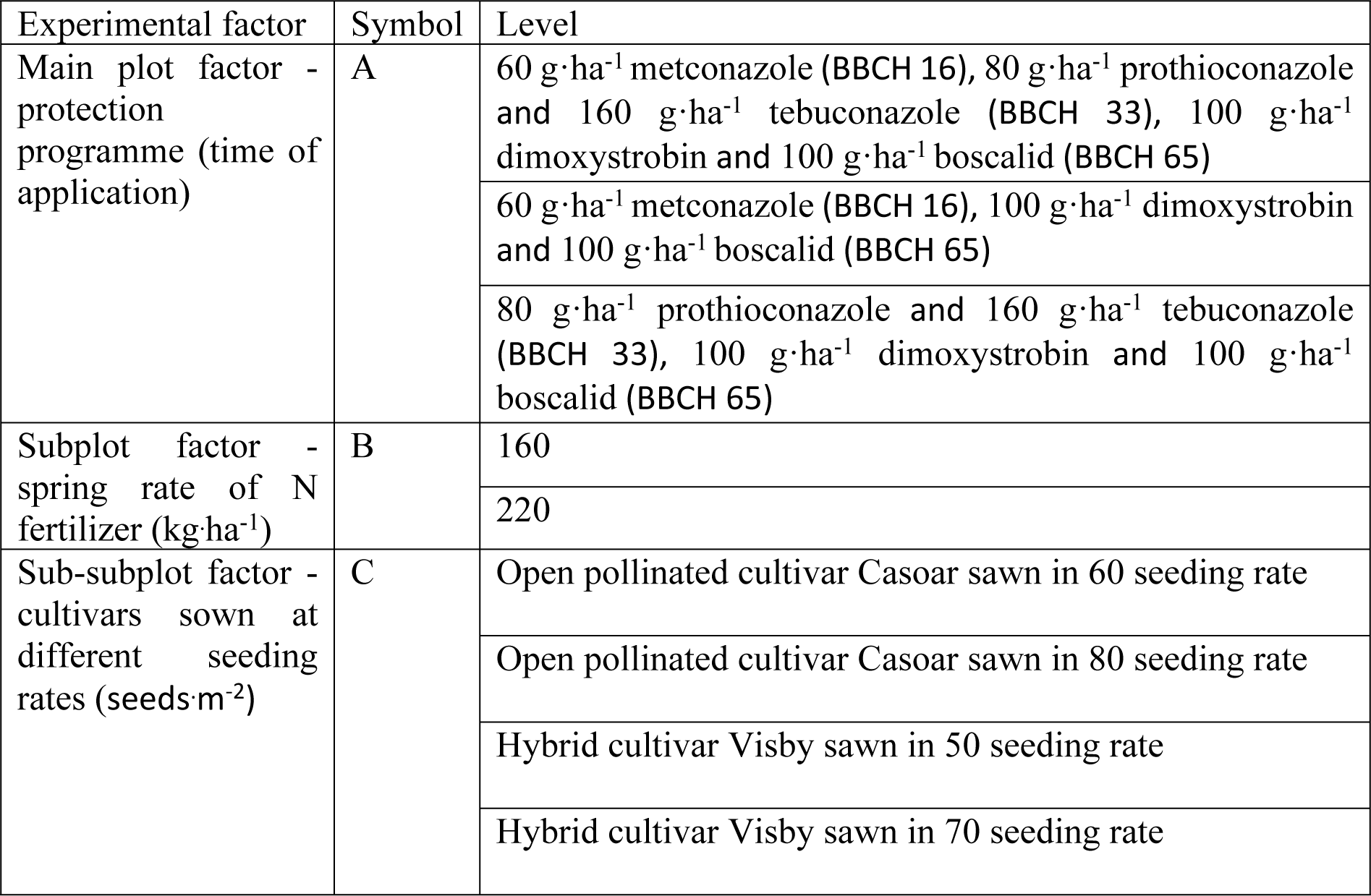
Experimental factors and levels in the experiment in split-split plot design

Disease identification and assessment of infection were performed at the ripening stage, when 40–50% of siliques were ripe—BBCH 84 and 85. The percentage of plants showing symptoms of white stem rot caused by *Sclerotinia sclerotiorum* and Phoma stem canker caused by *Leptosphaeria* spp. (anamorph *Phoma lingam*) and the percentage of the surface of silique showing *Alternaria* spot caused by *Alternaria* spp. and grey mould caused by *Botrytis cinerea* were determined. The percentage of plants showing symptoms of disease infection was estimated as mean disease incidence (DI) using the formula:

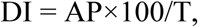

where AP is the number of affected plants and T is the total number of examined plants. Stem infection was assessed on all plants on a plot, and silique infection was estimated on 100 siliques randomly selected per plot. Disease severity of silique lesions was determined as the percentage of disease-affected area of each silique.

The plants were counted per unit area on each plot before and after winter and directly before harvest. Before the harvest, the number of siliques per plant were counted on 15 consecutive plants on each plot. Number of seeds per silique was determined by counting the number of seeds in 25 siliques sampled randomly from each plot. Weight of 1000 seeds was calculated from the weight of 400 seeds taken from the bulk yield samples. Based on the number of siliques per plant and the number of plants counted directly before harvest, the number of siliques per unit area was determined. Yield of seeds per plot was adjusted to 13% moisture content and then converted to Mg·ha^-1^.

The experimental data were compared using an analysis of variance (ANOVA). When *F*-ratio was significant, the least significant difference was calculated at *P*≤0.05 using Tukey’s test. ANOVA was performed using STATISTICA software (StatSoft Inc., 2011).

## Results

### Weather conditions and fenological crop development

The growing seasons in which the experiment was conducted differed significantly in meteorological conditions and influenced the fenological development of winter oilseed rape plants (Table 2). The length of the fall growing season ranged from 78 days in 2011 to 94 days in 2012, and total precipitation from 45.8 mm in 2011 to 109.9 mm in 2013. The most unfavourable weather condition in autumn was noted in 2013, when the shortage of precipitation before sowing, the cool and very rainy September and the cool first decade of October, and the stagnation of vegetation at the end of the second decade of November limited the development and the number of plants before winter. As a result of these conditions, oilseed rape plants developed only eight medium-sized leaves before winter, compared to 2011 and 2012 when the crop plants developed 12 large-sized leaves before winter and were in very good condition. The mean daily temperature during the winter dormancy was determined at −1.0 °C in 2012–2013 and at 2.9 °C in 2013–2014, while in the coldest decade of winter this parameter ranged from −12.4 °C in 2011–2012 to −5.3 °C in 2012–2013 (Table 2). In 2013 and 2014, the climatic condition during winter were conducive to crop overwintering, while in the coldest decade of 2012, frost reaching −20 °C at night, not accompanied by snowfall did not favour overwintering of oilseed rape plants. Due to the difference in the beginning of vegetation, the length of the spring growing season ranged from 106 days in 2013 to 122 days in 2014. The beginning of vegetation in spring was observed on 16 and 15 March in 2012 and 2014, respectively, while in 2013, on 8 April. Total precipitation was determined at 170 mm in 2012 and over 244 mm in 2014. A small amount of precipitation until the full flowering phase in 2012 did not favour the production of siliques. In 2013, a similar effect resulted from the late start of vegetation. Water conditions that were much more favourable for the development of siliques were observed in 2014. However, abundant rainfall during this flowering season was also conducive to plant infection by *S. sclerotiorum*. The seed harvest was carried out on 13, 23, and 15 July in 2012, 2013, and 2014, respectively.

**Table 2.** Fenological development of winter oilseed rape and weatcher conditions (2011-2014)

| Parameter | Period | Growing season |  |  |  |
| --- | --- | --- | --- | --- | --- |
|  |  | 2011/2012 | 2012/2013 | 2013/2014 | 1957-2014 |
| Number of days | Fall growth | 78 | 94 | 84 | 95 |
|  | Winter dormacy | 121 | 129 | 117 | 125 |
|  | Spring growth | 119 | 106 | 122 | 99 |
|  | flowering | 20 | 28 | 35 |  |
|  | Entire growing season | 318 | 329 | 323 | 319 |
| Total precipitation (mm)<br>*snowfall | Fall growth | 45.8 | 93.9 | 109.9 | 120.7 |
|  | The decade of sowing | 27.6 | 56.3 | 2.5 |  |
|  | September/first decade of October | 18.8/5.0 | 34.1/9.8 | 80.7/0.0 | 42.2/ |
|  | Winter dormacy | 143.3 | 171.5 | 69.6 | 133.4 |
|  | The coldest decade in winter | 1.0* | 17.3* | 20.5* |  |
|  | Spring growth | 170 | 213.0 | 244.4 | 200.0 |
|  | flowering | 22.8 | 48.6 | 81.7 |  |
|  | Entire growing season | 348.1 | 464.9 | 423.9 | 454.1 |
| Mean daily temperature (°C) | Fall growth | 9.6 | 9.4 | 9.5 | 8.8 |
|  | September/first decade of October | 15.2/12.9 | 14.5/10.4 | 12.6/8.4 | 13.6/ |
|  | Winter dormacy | 1.0 | -1.0 | 2.9 | 0.3 |
|  | The coldest decade in winter | -12.4 | -5.3 | -7.9 |  |
|  | Spring growth | 13.8 | 15.8 | 13.7 | 14.1 |
|  | flowering | 14.7 | 14.0 | 12.4 |  |
|  | Entire growing season | 7.9 | 7.2 | 8.6 | 7.7 |

### Disease control

The protective programs significantly affected the occurance of pathogens on the crop (Tabela 3). All effectively limited the disease symptoms (Table 4). Moreover, by affecting overwintering of plants and 1000 seed weight significantly affected the yield (Table 5). The average yield of seeds from the plants subjected to protective programmes was significantly higher than that from the unprotected plants by 420–560 kg·ha^-1^ (Table 6). The effectiveness of plant protection programmes depended on the environmental conditions that shaped the development of plants (Table 7). In the season 2011–2012, significantly higher yields were obtained on plots where plants were protected in autumn compared to the yields on control plots. The autumn treatment of plants with a fungicide with growth-regulating properties contributed to the limitation of plant losses during the winter which resulted in grater number of siliques per m^2^ compared to unprotected plots in autumn. However, in the season 2013– 2014, when plant development in the early stages was limited by the precipitation shortage before sowing and relatively low temperatures in September and the first decade of October (Table 2), significantly higher yields were obtained from the plots without the autumn treatment compared to the yields on the control plots. In the season 2012–2013, all methods of protection against pathogens significantly prevented the loss of yield. The yield from the protected plots was significantly higher than the yield from the unprotected plots by 420–580 kg·ha^-1^. None of the remaining experimental factors—cultivars sown at different seeding rates and levels of nitrogen fertilisation—had influence on the yield-protecting effect of fungicide treatment (Table 5).

**Table 3.** Anova *F*-test statistics

| Effect | *S.*<br>*sclerotiorum* | *Leptosphaeria*<br>*spp.* | *B. fuckeliana* | *Alternaria* spp. |
| --- | --- | --- | --- | --- |
|  | Disease incidence (%) |  | Disease severity (% of disease-affected area of each silique) |  |
| Growing season (Y) | 203.75** | 3.50* | 2.21* | 279.98** |
| Protection programme (time of application) (A) | 284.44** | 11.91** | 2.85* | 25.37** |
| Spring rate of N fertilizer (kg·ha<sup>-1</sup>) (B) | 0.08 ns | 0.22 ns | 0.00 ns | 2.24 ns |
| Cultivars sown at different seeding rates (C) | 3.69* | 6.09** | 1.14 ns | 1.16 ns |
| YXA | 2.11 ns | 2.16 ns | 1.96 ns | 2.45 ns |
| YXB | 0.27 ns | 0.54 ns | 0.43 ns | 0.11 ns |
| YXC | 1.66 ns | 1.83 ns | 1.96 ns | 1.45 ns |
| AXB | 0.10 ns | 0.68 ns | 0.10 ns | 1.82 ns |
| AXC | 1.11 ns | 1.09 ns | 1.02 ns | 1.14 ns |
| BXC | 0.82 ns | 0.77 ns | 0.90 ns | 0.34 ns |
| YXAXB | 0.75 ns | 0.70 ns | 0.53 ns | 0.59 ns |
| YXAXBXC | 1.05 ns | 0.40 ns | 1.06 ns | 0.41 ns |

ns-not significant
\*Significant P<0.05
\*\*Significant P<0.01

**Table 4.** Significance of differences between mean values of evalueted factors in evaluation of disease symptoms on plants cauosed by parhogens

| Factor/level | *S. sclerotiorum* | *Leptosphaeria* spp. | *B. fuckeliana* | *Alternaria* spp. |
| --- | --- | --- | --- | --- |
|  | Disease incidence (%) |  | Disease severity (% of disease-affected area of each silique) |  |
| Growing season |  |  |  |  |
| 2012 | 4.2 b<sup>1</sup> | 14.0 a | 4.46 a | 13.9 a |
| 2013 | 8.0 b | 10.3 ab | 1.78 b | 3.2 b |
| 2014 | 6.1 ab | 7.6 b | 4.12 a | 4.1 b |
| Protection programme (time of application) |  |  |  |  |
| BBCH 16 + BBCH 33 + BBCH 65 | 1.9 b | 8.3 b | 3.01 b | 5.77 b |
| BBCH 16 + BBCH 33 | 4.0 b | 9.9 b | 3.10 b | 6.23 b |
| BBCH 33 + BBCH 65 | 2.3 b | 10.9 b | 3.14 b | 6.12 b |
| Control | 28.6 a | 13.4 a | 4.56 a | 10.1 a |
| Spring rate of N fertilizer (kg·ha<sup>-1</sup>) |  |  |  |  |
| 160 | 9.1 a | 10.8 a | 3.45 a | 6.8 a |
| 220 | 9.2 a | 10.5 a | 3.45 a | 7.4 a |
| Cultivars sown at different seeding rates |  |  |  |  |
| Casoar 60 | 9.9 a | 8.4 b | 3.10 a | 6.9 a |
| Casoar 80 | 9.9 a | 10.3 ab | 3.83 a | 6.6 a |
| Visby 50 | 8.2 b | 11.4 ab | 3.29 a | 7.2 a |
| Visby 70 | 8.8 ab | 12.5 a | 3.61 a | 7.6 a |

<sup>1</sup> Means with the same letter are not significantly different at P<0.05 according to Tukey's HSD test

**Table 5.** Anova *F*-test statistics

| Effect | Seed yield<br>(Mg·ha<sup>-1</sup>) | Plants ·m<sup>-2</sup> |  | Overwintering (%) | Plants ·m<sup>-2</sup><br>before<br>harvest | Silique·plant<sup>-1</sup> | Silique·m<sup>-2</sup> | Seeds·silique<sup>-1</sup> | 1000 seed<br>weight (g) |
| --- | --- | --- | --- | --- | --- | --- | --- | --- | --- |
|  |  | before<br>winter | after<br>winter |  |  |  |  |  |  |
| Growing<br>season (Y) | 650.03** | 25.94** | 84.5** | 256.80** | 90.90** | 154.68** | 281.92** | 9.71* | 23.36** |
| Protection<br>programme<br>(time of<br>application)<br>(A) | 25.16** | 0.57 ns | 1.34ns | 2.92* | 1.58 ns | 2.88 ns | 4.47* | 2.43 ns | 31.92** |
| Spring rate of<br>N fertilizer<br>(kg·ha<sup>-1</sup>) (B) | 5.70** | 1.46 ns | 0.84 ns | 0.00 ns | 0.94 ns | 0.54 ns | 4.50* | 0.01 ns | 1.76 ns |
| Cultivars<br>sown at<br>different<br>seeding rates<br>(C) | 140.61** | 43.43** | 35.09** | 71.27** | 34.71** | 7.73** | 11.26** | 161.51** | 138.96** |
| YXA | 3.93** | 0.95 ns | 0.84 ns | 4.64 ** | 0.76 ns | 0.99 ns | 2.58* | 1.93 ns | 11.16** |
| YXB | 3.12* | 2.24 ns | 2.10 ns | 0.03 ns | 2.08 ns | 1.16 ns | 0.43 ns | 1.01 ns | 0.99 ns |
| YXC | 37.35** | 3.09** | 55.67** | 60.11** | 48.63** | 0.37 ns | 8.04 ** | 0.26 ns | 0.74 ns |
| AXB | 1.07 ns | 0.36 ns | 0.76 ns | 1.57 ns | 0.80 ns | 0.97 ns | 0.68 ns | 0.88 ns | 0.96 ns |
| AXC | 1.29 ns | 1.27 ns | 1.54 ns | 1.85 ns | 1.77 ns | 0.28 ns | 1.73 ns | 0.88 ns | 0.52 ns |
| BXC | 0.59 ns | 0.67 ns | 1.53 ns | 0.63 ns | 2.05 ns | 0.28 ns | 0.34 ns | 1.33 ns | 0.99 ns |
| YXAXB | 1.90 ns | 0.85 ns | 1.20 ns | 2.61 ns | 0.99 ns | 1.42 ns | 0.91 ns | 0.75 ns | 0.82 ns |
| YXAXBXC | 0.60 ns | 1.09 ns | 1.19ns | 1.01 ns | 1.26 ns | 0.25 ns | 0.91 ns | 1.24 ns | 2.09 ns |

ns-not significant
\*Significant P<0.05
\*\*Significant P<0.01

**Table 6.** Significance of differences between mean values of evalueted factors in evaluation of seed yield. plant overwintering and yield components

| Factor/level | Seed yield (Mg·ha<sup>-1</sup>) | Plants ·m<sup>-2</sup> |  | Overwintering (%) | Plants ·m<sup>-2</sup> before harvest | Silique·plant<sup>-1</sup> | Silique·m<sup>-2</sup> | Seeds·silique<sup>-1</sup> | 1000 seed weight (g) |
| --- | --- | --- | --- | --- | --- | --- | --- | --- | --- |
|  |  | before winter | after winter |  |  |  |  |  |  |
| Growing season |  |  |  |  |  |  |  |  |  |
| 2011/2012 | 1.82 c<sup>1</sup> | 48.9 ab | 24.3 b | 53.2 b | 23.3 b | 99 b | 2143 c | 21.4 b | 5.48 a |
| 2012/2013 | 5.27 b | 57.2 a | 56.2 a | 98.2 a | 55.2 a | 91 b | 4795 b | 22.1 a | 5.08 b |
| 2013/2014 | 6.39 a | 30.2 b | 28.6 b | 95.2 a | 28.6 b | 236 a | 6111 a | 22.7 ab | 5.31 ab |
| Protection programme (time of application) |  |  |  |  |  |  |  |  |  |
| BBCH 16 + BBCH 33 + BBCH 65 | 4.63 a | 44.6 a | 36.2 a | 83.4 a | 36.1 a | 144 a | 4448 a | 22.4 a | 5.29 a |
| BBCH 16 + BBCH 33 | 4.73 a | 46.3 a | 39.3 a | 84.2 a | 37.6 a | 131 a | 4408 a | 22.6 a | 5.36 a |
| BBCH 33 + BBCH 65 | 4.59 a | 46.3 a | 35.5 a | 79.7 b | 34.5 a | 155 a | 4547 a | 21.5 a | 5.34 a |
| Control | 4.17 b | 44.5 a | 35.5 a | 81.4 b | 34.7 a | 136 a | 3995 b | 22.0 a | 5.17 b |
| Spring rate of N fertilizer (kg·ha<sup>-1</sup>) |  |  |  |  |  |  |  |  |  |
| 160 | 4.47 b | 44.8 a | 35.9 a | 82.2 a | 35.3 a | 140 a | 4202 b | 22.1 a | 5.28 a |
| 220 | 4.58 a | 46.0 a | 36.8 a | 82.2 a | 36.1 a | 144 a | 4498 a | 22.1 a | 5.30 a |
| Cultivars sown at different seeding rates |  |  |  |  |  |  |  |  |  |
| Casoar 60 | 3.90 c | 42.9 b | 31.2 b | 72.7 b | 31.0 b | 141 b | 3764 b | 18.6 b | 5.48 a |
| Casoar 80 | 4.31 b | 55.1 a | 39.0 a | 73.8 b | 38.9 a | 118 c | 3949 b | 19.0 b | 5.42 a |
| Visby 50 | 4.76 a | 37.0 c | 33.5 b | 91.5 a | 32.0 b | 166 a | 4409 a | 25.4 a | 5.14 b |
| Visby 70 | 5.04 a | 46.7 b | 41.7 a | 90.7 a | 40.5 a | 142 b | 4685 a | 25.3 a | 5.11 b |

<sup>1</sup> Means with the same letter are not significantly different at P<0.05 according to Tukey's HSD

**Tabela 7.** Significance of differences between mean values of evalueted factors of growing season and protection programme in evaluation of seed yield, plant overwintering and yield components

| Growing season | Protection programme (time of application) | Seed yield (Mg·ha<sup>-1</sup>) | Plants ·m<sup>-2</sup> |  | Overwintering (%) | Plants ·m<sup>-2</sup> before harvest | Silique·plant<sup>-1</sup> | Silique·m<sup>-2</sup> | Seeds·silique<sup>-1</sup> | 1000 seed weight (g) |
| --- | --- | --- | --- | --- | --- | --- | --- | --- | --- | --- |
|  |  |  | before winter | after winter |  |  |  |  |  |  |
| 2011/2012 | BBCH 16 + BBCH 33 + BBCH 65 | 2.09 ab<sup>1</sup> | 47.1 a | 25.3 a | 59.3 a | 26.1 a | 105 a | 2415 a | 21.6 a | 5.39 a |
|  | BBCH 16 + BBCH 33 | 2.18 a | 49.0 a | 27.6 a | 59.1 a | 26.3 a | 89 a | 2398 a | 22.1 a | 5.53 a |
|  | BBCH 33 + BBCH 65 | 1.74 bc | 52.8 a | 22.6 a | 45.7 b | 20.6 a | 109 a | 1951 b | 21.0 a | 5.50 a |
|  | Control | 1.66 c | 46.9 a | 21.8 a | 48.6 b | 20.4 a | 91 a | 1806 b | 20.9 a | 5.49 a |
| 2012/2013 | BBCH 16 + BBCH 33 + BBCH 65 | 5.41 a | 56.2 a | 55.2 a | 98.2 a | 54.2 a | 93 a | 4808 a | 21.5 a | 5.11 a |
|  | BBCH 16 + BBCH 33 | 5.47 a | 57.5 a | 56.5 a | 98.2 a | 55.5 a | 93 a | 4861 a | 22.8 a | 5.15 a |
|  | BBCH 33 + BBCH 65 | 5.31 a | 55.9 a | 54.9 a | 98.1 a | 53.9 a | 93 a | 4872 a | 22.3 a | 5.12 a |
|  | Control | 4.89 b | 59.1 a | 58.1 a | 98.1 a | 57.1 a | 84 a | 4639 a | 22.0 a | 4.96 a |
| 2013/2014 | BBCH 16 + BBCH 33 + BBCH 65 | 6.37 ab | 30.4 a | 28.1 a | 92.9 a | 28.1 a | 235 a | 6123 b | 23.9 a | 5.35 a |
|  | BBCH 16 + BBCH 33 | 6.54 ab | 32.4 a | 30.9 a | 95.4 a | 30.9 a | 212 a | 5963 b | 22.9 a | 5.41 a |
|  | BBCH 33 + BBCH 65 | 6.72 a | 30.2 a | 28.9 a | 95.2 a | 28.9 a | 262 a | 6818 a | 21.1 a | 5.41 a |
|  | Control | 5.95 b | 27.6 a | 26.6 a | 97.3 a | 26.6 a | 234 a | 5541 c | 23.0 a | 5.06 b |

### Rate of nitrogen fertilisation

The increase in the level of nitrogen fertiliser from 160 to 220 kg·ha^-1^ caused a significant increase in the average seed yield (Table 6). The effectiveness of nitrogen fertilisation varied during the years of the study. The nitrogen fertilisation significantly increased the seed yield in the second and the third season (2012–2013 and 2013–2014) (Table 8). The effectiveness of nitrogen fertilisation was not influenced neither by cultivars sown at different seeding rates nor by fungicide treatment (Table 5). The higher yield obtained with 220 kg N·ha^-1^ can be attributed to the higher number of siliques per unit area (Table 6). The number of siliques per unit area was the only yield component that significantly increased with an increase in the rate of nitrogen fertilisation (220 kg·ha^-1^). Similar to the yield, this parameter also significantly increased with the increase in the rate of nitrogen fertilisation in the second and the third experimental cycle (2012–2013 and 2013–2014) (Table 8).

**Table 8.** Significance of differences between mean values of evalueted factors of growing season and spring nitrogen fertilization in evaluation of seed yield, plant overwintering and yield components

| Growing season | Spring rate of N fertilizer (kg·ha<sup>-1</sup>) | Seed yield (Mg·ha<sup>-1</sup>) | Plants m<sup>-2</sup> |  | Overwintering (%) | Plants m<sup>-2</sup> before harvest | Silique plant<sup>-1</sup> | Silique m<sup>-2</sup> | Seeds siique<sup>-1</sup> | 1000 seed weight (g) |
| --- | --- | --- | --- | --- | --- | --- | --- | --- | --- | --- |
|  |  |  | before winter | after winter |  |  |  |  |  |  |
| 2011/2012 | 160 | 1.93 a<sup>1</sup> | 46.9 a | 22.7 a | 53.0 a | 21.8 a | 100 a | 2060 a | 21.6 a | 5.47 a |
|  | 220 | 1.90 a | 51.0 a | 25.9 a | 53.4 a | 24.9 a | 96 a | 2225 a | 21.2 a | 5.48 a |
| 2012/2013 | 160 | 5.15 b | 57.6 a | 56.6 a | 98.2 a | 55.6 a | 84 a | 4533 b | 22.2 a | 5.08 a |
|  | 220 | 5.39 a | 56.8 a | 55.8 a | 98.1 a | 54.8 a | 98 a | 5057 a | 22.1 a | 5.08 a |
| 2013/2014 | 160 | 6.34 b | 30.0 a | 28.6 a | 95.3 a | 28.6 a | 235 a | 6012 b | 22.4 a | 5.28 a |
|  | 220 | 6.45 a | 30.3 a | 28.7 a | 95.1 a | 28.7 a | 237 a | 6210 a | 23.1 a | 5.34 a |

<sup>1</sup> Means with the same letter are not significantly different at P<0.05 according to Tukey's HSD

### Cultivars sown at different seeding rates

The average yield in the 3-year experiment ranged from 1.82 to 6.39 Mg·ha^-1^ (Table 6). The highest seed yield was observed in the season with favourable wintering and water conditions for the stages of flowering and fruit growth (2013–2014), whereas the lowest yield was noticed in the year with the least favourable conditions for dormancy (2011–2012). Regardless of the seeding rate, the hybrid cultivar Visby showed the highest average yields (5.04 Mg·ha^-1^ at 70 seeds·m^-2^ and 4.76 Mg·ha^-1^ at 50 seeds·m^-2^) in the study. The average yield of the open-pollinated cultivar Casoar was more dependent on the seeding rate; a higher yield (4.31 Mg·ha^-1^) was obtained at the higher seeding rate (80 seeds·m^-2^), whereas lowering the seeding rate to 60 seeds·m^-2^ resulted in a yield decrease by 410 kg·ha^-1^. The effect of seeding rate on the yield of the assessed cultivars was dependant on the weather conditions during the seasons in which the experiment was conducted (Table 9). In the least favourable conditions for plant development (2011–2012), significant differences in the yields were noted in Visby cultivar sown at different seeding rates (2.87 Mg·ha^-1^ at 70 seeds·m^-2^ and 2.29 Mg·ha^-1^ at 50 seeds·m^-2^). However, in the most favourable conditions (2013–2014), significant differences were observed in the yields of Casoar cultivar (6.54 Mg·ha^-1^ at 80 seeds·m^-2^ and 5.82 Mg·ha^-1^ at 60 seeds·m^-2^). Moreover, in these conditions, and also in the conditions of the 2012–2013 season, the yields of Casoar cultivar from the plots sown at a higher seeding rate (80 seeds·m^-2^) did not differ significantly from the yields of Visby cultivar from the plots sown at 50 and 70 seeds·m^-2^.

**Table 9.** Significance of differences between mean values of evalueted factors of growing season and cultivar sawn in seeding rate in evaluation of seed yield, plant overwintering and yield components

| Growing season | Cultivars sown at different seeding rates | Seed yield (Mg·ha<sup>-1</sup>) | Plants ·m<sup>-2</sup> |  | Overwintering (%) | Plants ·m<sup>-2</sup> before harvest | Silique·plant<sup>-1</sup> | Silique·m<sup>-2</sup> | Seeds·silique<sup>-1</sup> | 1000 seed weight (g) |
| --- | --- | --- | --- | --- | --- | --- | --- | --- | --- | --- |
|  |  |  | before winter | after winter |  |  |  |  |  |  |
| 2012 | Casoar 60 | 1.20 c<sup>1</sup> | 42.9 b | 12.0 b | 29.3 b | 12.3 b | 101 a | 1034 b | 18.9 b | 5.69 a |
|  | Casoar 80 | 1.02 c | 59.8 a | 14.0 b | 26.8 b | 14.6 b | 76 a | 1148 b | 18.7 b | 5.62 a |
|  | Visby 50 | 2.29 b | 40.9 b | 32.3 a | 81.1 a | 30.1 a | 113 a | 3296 a | 23.9 a | 5.31 b |
|  | Visby 70 | 2.87 a | 52.3 ab | 38.9 a | 75.7 a | 36.3 a | 92 a | 3082 a | 24.0 a | 5.27 b |
| 2013 | Casoar 60 | 4.87 b | 53.3 b | 52.2 bc | 98.0 a | 51.2 bc | 92 a | 4483 a | 18.7 b | 5.25 a |
|  | Casoar 80 | 5.20 ab | 69.5 a | 68.4 a | 98.5 a | 67.5 a | 69 a | 4581 a | 19.7 b | 5.22 a |
|  | Visby 50 | 5.48 a | 47.4 b | 46.4 c | 97.9 a | 45.4 c | 110 a | 4971 a | 25.4 a | 4.94 b |
|  | Visby 70 | 5.53 a | 58.6 ab | 57.6 b | 98.2 a | 56.6 b | 95 a | 5146 a | 24.9 a | 4.92 b |
| 2014 | Casoar 60 | 5.82 b | 32.6 ab | 29.4 ab | 90.8 a | 29.4 ab | 231 a | 5998 a | 18.3 b | 5.49 a |
|  | Casoar 80 | 6.54 a | 36.1 a | 34.7 a | 96.2 a | 34.7 a | 205 a | 6385 a | 18.6 b | 5.43 a |
|  | Visby 50 | 6.50 a | 22.7 b | 21.7 b | 95.6 a | 21.7 b | 270 a | 5419 a | 26.9 a | 5.17 b |
|  | Visby 70 | 6.72 a | 29.3 ab | 28.6 ab | 98.2 a | 28.6 ab | 235 a | 6642 a | 27.2 a | 5.15 b |

Higher yields of Visby cultivar can be attributed to the higher number of seeds per silique and higher number of siliques per m^2^ resulting from the higher number of plants per m^2^ before harvest despite the lower seeding rate of 10 seeds·m^-2^ and the greater compensation ability expressed by the greater number of siliques per plant (Table 6). Average higher number of Visby cultivar plants before harvest observed with the lowest seeding rate resulted from the greater overwintering success of this cultivar in the season with the worst conditions of dormancy. In turn, the higher number of siliques per plant of Visby cultivar was a result of the higher plant height. On average, a Visby cultivar plant was taller than a Casoar cultivar plant by 15 cm (data not shown). Irrespective of the cultivar, an increase in seeding rate (Visby: 50– 70 pure live seeds·m^-2^; Casoar: 60–80 pure live seeds·m^-2^) decreased the number of siliques per plant and the weight of 1000 seeds. However, significant differences were recorded between the cultivars only in the number of siliques per plant.

## Discussion

### Disease control

The winter oilseed rape is exposed to disease infections throughout the growing season. The loss of yields is caused, on the one hand, by infection depending on the genetically controlled resistance of cultivars [38] and the effectiveness of protection from pathogens, and on the other hand, by the incidence of pathogens that cause the most dangerous diseases in this species, such as blackleg (*P. lingam*, syn. *Leptosphaeria maculans*), stem rot (*S. sclerotiorum*), light leaf spot (*Cylindrosporium concentricum*, syn. *Pyrenopeziza brassicae*), verticillium wilt (*Verticillium dahliae*), dark pod spot (*Alternaria brassicae*), downy mildew (*Peronospora parasitica*), grey mould (*B. cinerea*), and clubroot disease (*Plasmodiophora brassicae*) [31, 42, 48], triggered by the environmental and agrotechnical conditions. The conducted experiment showed that the applied fungicides limited the yield losses resulting from pathogen infection and unfavourable wintering conditions (Table 4, 6, 7). The presented results thus broaden the view of Kruse and Verreet [21] that the increase in yield due to fungicides treatment is a result of not only the inhibition of infection by pathogens but also the action of fungicides as a growth regulator, which contributes to shortening the main shoot, reducing plant lodging, and increasing the resistance of siliques to cracking. In the season 2011–2012 characterised by severe and snowless winter, the use of chemical fungicide with growth-regulating properties at the six-leaf phase (BBCH 16) allowed limiting the loss of plants during winter dormancy and thereby resulting in higher crop yields from the protected plots in autumn (Tbla 7). This result is also confirmed by the previous studies of Geisler [11], Paul [27], and Schulz [34], which showed an increase in the winter hardiness of winter oilseed rape as a result of shortening the plant growth with the use of growth regulators. However, in the season 2013–2014, autumn treatment was not very effective (Table 7). The winter was mild (Table 2), and the treatment at the six-leaf stage of winter oilseed rape plants that were poorly developed due to unfavourable weather conditions did not significantly reduce the yield losses (Table 7). During this season, significantly higher yields were obtained with spring treatment, compared to the yields collected from control plots. Although the effectiveness of all the applied protective methods was significant only in the 2012–2013 season, it is worth emphasising that during the experiment period, the lowest yields were always obtained from the unprotected plots. This indicates the effectiveness of the chemical protection programme used in the experiment in reducing plant infection. The programme that included treatment at the full flowering stage (BBCH 65) was the most effective in reducing the symptoms of stem rot (*S. sclerotiorum*) and dark pod spot (*A. brassicae*) (Table 5). The effectiveness of the treatment in the phase of full flowering was also confirmed by Jankowski et al. [17] and Kruse and Verreet [21]. In turn, the combination of the autumn and early-spring treatments was the most effective in limiting the disease symptoms caused by *P. lingam*, syn. *L. maculans* (Table 5). Similar results were also reported by Kruse and Verreet [21]. Moreover, the experiment did not show any differences in yield between the cultivars as a result of intensification of protection, confirming the earlier reports of Jajor et al. [15], Jędryczka and Kaczmarek [18], and Wójtowicz [48] and thus indirectly indicating the need to include a chemical protection programme in the cultivation of winter oilseed rape.

### Rate of nitrogen fertilisation

Spring nitrogen fertilisation is considered to be one of the most important factors of production [4, 31]. In many experiments, yield increase has been achieved within the dose limits of 150– 180 kg N·ha^-1^ [3, 4, 35], and therefore, high (about 240 kg) [46, 50] and very high doses of nitrogen fertilisation (about 300 kg) [36] are rarely required. Determination of the optimal nitrogen dose is hampered by the strong dependence of fertilisation efficiency on the changing weather conditions in the years. In the present work, the increase in the level of fertilisation from 160 to 220 kg·ha^-1^ proved to be effective in the second and third growing seasons (2012– 2013, 2013–2014). However, in the first season (2011/2012), in the conditions of severe and snowless winter and shortage of rainfall during spring development (Table 2), the increase in the level of fertilisation was ineffective (Table 8). The obtained results correspond with the results presented by Jankowski et al. [17], which also showed the variability of fertilisation efficiency in the years of investigations, when in one season there was a significant increase in the yield at a dose of 240 kg·ha^-1^, and in another at a dose of 180 kg·ha^-1^. The present research (Table 5) also confirms the results of studies describing a similar response of cultivars to nitrogen fertilisation [8, 17], despite their diverse ability to take up and use nitrogen [19, 44]. Therefore, unequal nitrogen uptake and utilisation abilities are usually not significant enough to contribute to a significant difference in crop yield that can result from the varied level of nitrogen fertilisation in the range of doses recommended for agricultural practice. An indirect confirmation of the above statement is the small number of scientific reports showing a significantly different response of cultivars to nitrogen fertilisation. Howewer Wielebski and Wójtowicz [43] showed a significantly lower dependence of the yield level on the level of nitrogen fertilisation of the first hybrid cultivar Synergy in comparison with population cultivars. A similar tendency was confirmed by the research of Pellet [28], except that no statistically significant differences were shown by the results. The results of the present study (Table 4) are also in line with the results presented by Sadowski et al.[33] and Lemańczyk et al. [23], which showed that the increase in nitrogen fertilisation did not increase the severity of disease symptoms caused by *S. sclerotiorum* and *Leptosphaeria* ssp. The lack of a negative impact of increased nitrogen fertilisation on the infection of plants by the most dangerous pathogens of oilseed rape is desirable from the point of view of production intensification.

### Cultivars sown at different seeding rates

The yield level is an indicator of a plant’s development which depends on its response to environmental and agrotechnical conditions. The plant response is in turn mainly conditioned by the impact of environmental and agrotechnical factors and its adaptability to adverse conditions. In the unfavourable seasons, yields are significantly determined by the plant resistance to stress. Compared to the Casoar cultivar, over 1-tonne higher yield was recorded for the Visby cultivar in the 2011–2012 season (Table 9), characterised by severe and snowless winter (Table 2), due to the greater winter hardiness and consequently the better wintering of this cultivar, which resulted in a much greater number of siliques per unit area (by more than 1000 per m^2^). In the conditions of severe and snowless winter and shortage of rainfall during spring development in the season 2011–2012, the yield of the Visby cultivar was influenced mainly by the derivative of the sowing rate—the number of plants per unit area—as evidenced by the higher yields (over 0.5 tonne) collected from the plants sown densely at 70 seeds·m^-2^ (Table 9). In the season 2013–2014 characterised by a shortage of rainfall during emergence and good moisture content in spring, the amount of seeds sown determined the yields of the Casoar cultivar which showed a significant difference. Significantly higher yields were collected from the Casoar cultivar plants sown more densely at 80 seeds·m^-2^. The lack of a significant variation in the yield of the hybrid cultivar in this season indicates that due to its greater vigour it was able to better utilise the favourable humidity conditions recorded in spring 2014. In the remaining growing seasons, the variation in the yield level between the plots sown at different rates was statistically insignificant. Nevertheless, in the case of both hybrid and open-pollinated cultivars, higher yields were collected from the plants sown more densely. These results are consistent with the study by Jankowski et al. [17], which assessed the impact of the seeding rate (80, 60, and 40 seeds·m^-2^) of hybrid cultivars on their yield and showed that the highest yields were obtained from densely sown plants (80 seeds·m^-2^). Similar results are found in the work of Wójtowicz et al. [49], which revealed a significant reduction in the yield of the hybrid cultivar when the amount of seeds sown was reduced from 70 to 35 seeds·m^-2^. Experiments showed that in the conditions of Wielkopolska higher yields were obtained at higher seeding rates as a consequence of the unfavourable humidity conditions during emergence and thermal conditions during the winter dormancy period resulting in a reduction in the number of plants per unit area and the late beginning of vegetation and shortage of precipitation in the spring limiting the production of siliques. This broadens the view presented by Jankowski and Budzyński [16] about the influence of thermal conditions during winter and humidity in spring on the yields from plots sown at varied seeding rates. The humidity conditions during emergence also play an equally important role and can significantly adversely affect the number of plants per unit area. Earlier results of Wójtowicz et al. [49] proved that adverse conditions causing a decrease in plant density can occur with high probability during plant emergence.

In the three years of experiment (2009–2011) conducted in Wielkopolska region, unfavourable conditions during early autumn development, which contributed to a reduction in the number of plants in relation to the amount of seeds sown by about 40%, were recorded in two growing seasons (2009–2010 and 2010–2011). From the presented results (Table 5), the lack of a significant interaction between the amount of seeds sown for the evaluated cultivars and the applied levels of nitrogen fertilisation is also worth noting. The above dependence suggests that in the experimental conditions higher plant density did not limit plant development. These results are consistent with those of Budzyński [4], who in intensive technologies for excessive compaction for hybrid cultivars in spring recognised 60 plants·m^-2^. Despite the documented possibility of reducing the amount of seeds sown to 20–40 pure live seeds·m^-2^ [20, 22, 40], the presented results force taking into account the humidity conditions when determining the seeding rate, especially in the areas characterised by higher probability of precipitation shortage. In addition, the results lead to a hypothesis that under the conditions of predicted global warming, which will result in greater weather variability, the role of the amount of seeds sown, a basic element of rapeseed agrotechnology, in yielding will increase. Another factor that will be of more importance in the future is the selection of cultivar for cultivation. Furthermore, the more frequent occurrence of unfavourable conditions for the development of crop plants will require looking for stable-yielding cultivars.

## Conclusion

The conducted experiment showed that all the analysed factors had a significant impact on the yield level. The applied fungicides limited the crop losses resulting from pathogen infection and unfavourable wintering conditions. The protection programme consisting of treatment at the BBCH 65 flowering stage most effectively reduced the damage caused by stem rot (*S. sclerotiorum*) and dark pod spot (*A. brassicae*). In turn, the protection programme combining the autumn and early-spring treatments most effectively limited the infection caused by *P. lingam*, syn. *L. maculans*. In addition, the effectiveness of nitrogen fertilisation varied during the years of the study. In the conditions of shortage of precipitation during spring development after a period of stress caused by severe and snowless winter, the increase in the rate of fertilisation up to 220 kg·ha^-1^ was ineffective. In less stressful conditions, nitrogen fertilisation exerted a yield-increasing effect up to a rate of 220 kg·ha^-1^. Increase in nitrogen fertilisation level also did not increase the severity of disease symptoms caused by *S. sclerotiorum* and *Leptosphaeria* ssp. This lack of a negative impact of increased nitrogen fertilisation on plant infection by the most dangerous rape pathogens is desirable from the point of view of production intensification. The yield of seeds depended both on the yield potential of the cultivar and its ability to develop under stressful conditions. The experiment has shown a greater resistance of the restored hybrid cultivar Visby to the adverse thermal conditions. Good winter hardiness allowed obtaining relatively high yields of this cultivar in the conditions of severe and snowless winter. Moreover, due to its greater vigour than Casoar cultivar, Visby cultivar was able to better utilise the favourable humidity conditions that were recorded in spring 2014, and regardless of the amount of seeds sown, its yield was high. By contrast, the yields of Casoar were more dependent on the amount of seeds sown. Nevertheless, both cultivars yielded higher at a higher sowing rate. They also responded similarly to plant protection programmes and the rate of nitrogen fertilisation in spring.

